# Whole genome sequences of *Treponema pallidum* subsp. *endemicum* isolated from Cuban patients: the non-clonal character of isolates suggests a persistent human infection rather than a single outbreak

**DOI:** 10.1101/2021.10.13.464209

**Authors:** Eliška Vrbová, Angel A. Noda, Linda Grillová, Islay Rodríguez, Allyn Forsyth, Jan Oppelt, David Šmajs

**Affiliations:** Department of Biology, Faculty of Medicine, Masaryk University, Brno, Czech Republic; Department of Mycology-Bacteriology, Institute of Tropical Medicine “Pedro Kourí”, Havana, Cuba; GeneticPrime Dx, Inc., La Jolla, California, 92037, United States of America; San Diego State University, San Diego, California, United States of America; Department of Pathology and Laboratory Medicine, Perelman School of Medicine, University of Pennsylvania, United States of America

**Keywords:** *Treponema pallidum* subsp. *endemicum*, bejel, Cuba, non-clonal character, whole genome sequences.

## Abstract

Bejel (endemic syphilis) is a neglected non-venereal disease caused by *Treponema pallidum* subsp. *endemicum* (TEN). Although it is mostly present in hot, dry climates, a few cases have been found outside of these areas. The aim of this work was the sequencing and analysis of TEN isolates obtained from “syphilis patients” in Cuba, which is not considered an endemic area for bejel.

Genomes were obtained by pool segment genome sequencing or direct sequencing methods, and the bioinformatics analysis was performed according to an established pipeline.

We obtained four genomes with 100%, 81.7%, 52.6%, and 21.1% of broad coverage, respectively. The sequenced genomes revealed a non-clonal character, with nucleotide variability ranging between 0.2–10.3 nucleotide substitutions per 100 kbp among the TEN isolates. Nucleotide changes affected 27 genes, and the analysis of the completely sequenced genome also showed a recombination event between *tprC* and *tprI*, in TP0488 as well as in the intergenic region between TP0127–TP0129.

Despite limitations in the quality of samples affecting broad sequencing coverage, the determined non-clonal character of the isolates suggests a persistent infection in the Cuban population rather than a single outbreak caused by imported case.

**Author summary:** The incidence of venereal syphilis has greatly increased in the last years, however endemic syphilis (bejel) which have been considered as a disease restricted to dry arid areas such as the Sahel and the Middle East, remain as a neglected disease. In Cuba, which is a tropical country, several bejel cases were unexpectedly detected few years ago in “syphilis” patients with no records of travel abroad or sex with foreign partners. In this study, we explored the whole genome sequences from four of the Cuban *Treponema pallidum* subsp. *endemicum* (TEN) isolates and the substantial genetic diversity detected among them suggests a persistent infection of TEN within the human population rather than a single outbreak of a TEN isolate introduced from an area where it is typically endemic. This finding has significant implications on this neglected and also possibly tropical disease in terms of geographical/temporal distribution, and highlights the importance of keeping in mind neglected diseases in apparently non-endemic areas.

## Introduction

*Treponema pallidum* ssp. *endemicum* (TEN) is the causative agent of endemic syphilis (bejel), a neglected non-venereal disease that is mostly present in hot, dry areas of the world. TEN treponemes are highly related (99.7% identity at the genome level) to the *Treponema pallidum* ssp. *pallidum* (TPA), the causative agent of syphilis [1].

Acute bejel infections are mostly found among children between two and 15 years. Like syphilis, bejel can be divided into disease stages. In the primary stage, a small, painless ulcer is usually found in the oral cavity or nasopharynx [2] and usually remains undetected. In the secondary stage, numerous lesions appear in several body areas. In the last stage, gummatous lesions or bone alterations can appear. In several documented cases, bejel treponemes also infected the nipples of nursing women or genital areas [3]. Transmission of this disease occurs typically through direct mucosal and skin contact or is transferred by eating utensils or drinking vessels [2].

While bejel is a typical disease in dry arid areas such as the Sahel and the Middle East [2], it has also been found in Canada [4], France [5], Japan [6], and Cuba [7]. Cases in Canada and France were explained by bejel being imported from endemic areas, Senegal and Pakistan, respectively. On the other hand, the bejel cases in Japan and Cuba were originally identified in patients suspected of having syphilis and with no evidence of disease import and having a sexual route of transmission.

In bacteriology, the strict definition of a clone tends to be loosened slightly, and clones are defined pragmatically as isolates that are indistinguishable or highly similar, using a particular molecular typing procedure [8]. Among TPA, certain predominant genotypes are observed to infect the human population, i.e., the allelic profile 1.3.1. according to the MLST system [9] and 14d/f using enhanced CDC molecular typing [10].

In the case of TEN, there are very few genomic analyses, and the two available whole genome sequences, differing in 37 single nucleotide variants, come from the reference strains Bosnia A [11] (CP007548) and Iraq B [12] (CP032303). In addition, nine recombinant loci have been detected in TEN isolates during typing studies [13].

In this work, we determined whole genome sequences of four TEN isolates from Cuba, one of which was complete. While previous molecular typing of bejel samples discovered in Cuba showed that they have a clonal character at the tested chromosomal loci, our work demonstrates that the four characterized TEN isolates have a non-clonal character that suggests the presence of long-term human infection rather than a single outbreak due to a specific introduction of TEN into the adult population.

## Methods

### Study design and clinical samples

This observational descriptive study includes four clinical samples from bejel patients collected in years 2014, 2015 and 2017. Patients attended to the Instituto de Medicina Tropical “Pedro Kourí,” Havana, Cuba and preliminary characteristics were previously published [7,14]. DNA was isolated using QIAmp DNA mini kits (Qiagen, Hilden, Germany) according to the manufacturer’s instructions. Following DNA isolation, whole genome amplification was performed using REPLI-g Single Cell kits (Qiagen, Hilden, Germany) [15]. The number of TEN copies in samples was determined by real-time PCR using primers targeting *pol*A [16]. Samples for sequencing were selected according to the (1) total volume of available sample, (2) number of DNA copies in the sample, and (3) percent positivity of PCR amplification of 14 different *T. pallidum* intervals [15].

### Pooled segment genome sequencing (PSGS)

Genomes of samples C178 and C279 were chosen for the PSGS method [11,15,17], reflecting successful ratios in the amplification of 14 TPIs of pool 1 (11 of 14 for C279 and six for C178). All primers are listed in Supplementary Table 1 [11]. PCR products were purified using QIAquick PCR Purification Kits (QIAGEN, Hilden, Germany) and were divided into four pools to separate paralogous genes; this was to avoid later misassembly of these genes. TPI 11A containing *tprC* from pool 1 and 25B containing *tpr*E from pool 2 was added to pool 4. For sample 178, all PCR products of TPI were mixed in one pool since there were no paralogous regions amplified. PCR products of individual pools were mixed in equimolar amounts. During construction of the DNA library using a Nextera ® XT DNA Sample Preparation Guide kit, these four pools were labeled with multiplex identifier (MID) adapters. Prepared pools were sequenced using NGS sequencing on an Illumina platform. The numbers of repetitions in the *arp* and TP0470 genes in sample C279 and the sequence of TP0488 in sample C178 were determined using Sanger sequencing.

Reads were aligned to references using Lasergene software (DNASTAR, Madison, WI, USA). Final sequences were assembled from the consensus of individual pools and Sanger sequenced regions.

### Direct sequencing

Samples C75 and C77 were directly sequenced as described previously [15]. Sample C77 was sequenced with and without DNA enrichment, and data from both approaches were combined in the analysis. For *Dpn*I enrichment, 10–40 μl of the clinical sample was mixed with *Dpn*I-coated beads to a final volume of 50 μl in 1.7 mL Eppendorf tubes as previously described [18].

The bioinformatic analysis was performed according to the pipeline described previously [19]. The quality of raw reads was checked using FastQC (v0.11.5) [20]; pre-processing used Cutadapt (v1.15) [21] and Fastx-toolkits (v0.0.14) [22]. The pre-processed reads were mapped to the human genome reference (hg38), and the human-matching reads were removed using BBMap (v37.25) [23]. The remaining reads were mapped to the TEN reference genome of Bosnia A (CP007548) using BWA MEM (v0.7.15) [24]. The post-processing of the mapping was performed using Samtools (v1.4) [25], Picard (v2.8.1) [26], GATK (v3.7) [27], and NGSUtils/bamutils (v0.5.9, commit a7f08f5) [28].

### Phylogenetic analyses

For phylogenetic analyses, draft genomes and a whole-genome of isolate C279 were compared with TEN strains Bosnia A (CP007548), Iraq B (CP032303), isolate 11q/j (KY120774-KY120814), and with Japanese bejel isolates Osaka-2014 (LC383799, LC430604), Kyoto-2017 (LC430601, LC430606), Osaka-2017A (LC383801, LC430605), Osaka-2017B (LC430602, LC430607), and Osaka-2018 (LC430603, LC430608). Phylogenetic trees were constructed using the Maximum Likelihood Method [29] with bootstrapping [30] in MEGA7 software [31].

### Annotation of Complete Genomes

For gene annotation, Geneious software (2021.1.1.) was used. The *tprK* gene showed intra-strain variability in all samples, and the corresponding nucleotide positions were denoted as “N.” Genes from the TEN C279 strain were tagged with the TENDC279_ prefix. In C279, the original locus tag numbering corresponds to the tag numbering of orthologous genes annotated in the TEN Bosnia A genome. In the draft genomes of C75, C77, and C178, and where it was possible, genes were annotated based on TEN Bosnia A tagged TENDC75, TENDC77, and TENDC178, respectively. Sequences from isolates C279 and C77 can be found under the following Accession Numbers: CP078090, CP081507.

### Ethics statement

The study protocol was approved by the Research Ethics Committee of IPK, and it was conducted in compliance with the Declaration of Helsinki.

## Results

Descriptions of the analyzed samples are shown in Supplementary Table 2, and the overall results of the whole genome sequencing are shown in Supplementary Table 3 and Supplementary Fig.

### Sequence diversity in the Cuban TEN genomes

We compared the fully determined whole genome sequence of TEN C279 with the partially determined genome sequences of C77, C75, and C178. Pairwise nucleotide diversity values determined among the corresponding genomes are presented in the Table and Fig 1A/B. The phylogenetic relationship among C279 and previously reported TEN isolates is presented in Fig 1C.

**Figure 1.**
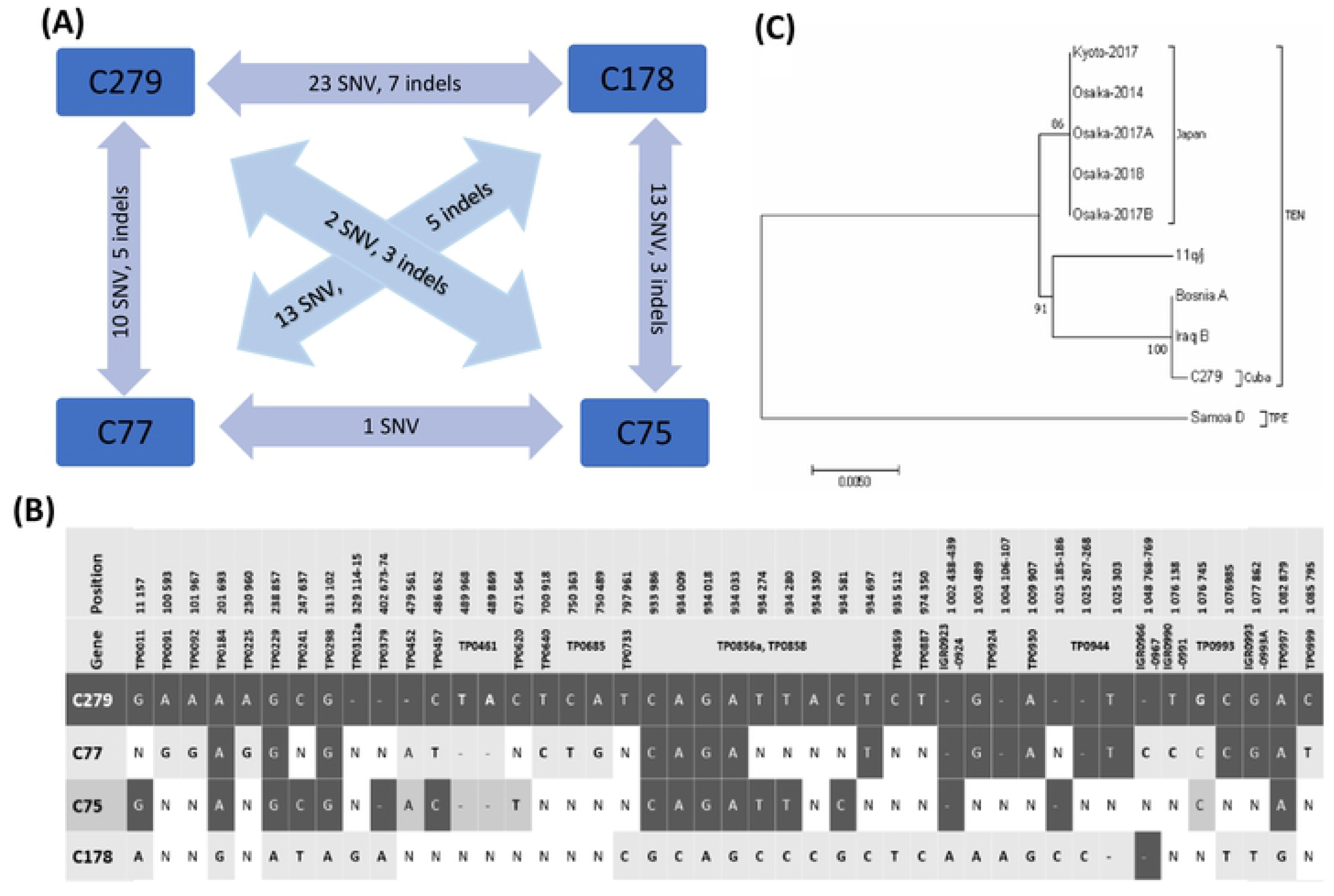
Genetic diversity among Cuban treponemal isolates. **(A)** The number of detected genetic differences among Cuban bejel strains (single nucleotide differences, iodels). **(B)** Detailed visualization of genetic differences runong Cuban bejel strains. N, the con esponding nucleotide was not determined. **(C)** Phylogenetic analysis of available TEN genomes or sequences. The evolutionary histo1y was inferred using the Maxi111u1n Likelihood method based on the Tamura-Nei model [41S]. The tree is drawn to scale, with branch lengths 111easured in the nu111ber of substitutions per site. The percentage of trees in which the associated taxa clustered together is shown next to the branches. For the construction of the tree, regions TP0548 (between coordinates 591,226-592,285) and TP0856 (between coordinates 932,947 - 933,I 82) [4] were used. C75, C77, and Cl78 were omitted due to unavailable sequences. The tree with the highest log likelihood (−1994.76) is shown. The analysis involved IO nucleotide sequences. There were a total of 1175 positions in the final dataset. As a root, TPE strain Samoa D (CP002374) was used.

Compared to the complete genome of TEN Bosnia A, TEN Cuban genomes showed 3.9–39.4 nucleotide substitutions per 100 kb of the genome, varying by one order of magnitude. The most divergent genomes were C279 and C178, with a nucleotide diversity of 39.4 and 20.4, respectively. On the other hand, diversity from Bosnia A was relatively small in samples C75 and C77. Genetic diversity within the Cuban samples was in the range 0.2–10.3, i.e., somewhat lower compared to the diversity between Cuban samples and Bosnia A.

### New recombination events in the C279 genome

In addition to the inter-strain recombination events at the TP0488 and TP0548 loci described earlier [5]; the C279 genome contains (compared to Bosnia A) a total of 332 changes in the *tprC* gene (TP0117), which is, in part, sequentially identical to the *tprI* gene of Bosnia A (Fig 2). The *tprC* of C279 is thus a result of an intrastrain recombination event where the recombinant gene contains about half of *tprI* copied to *tprC.*

**Figure 2.**
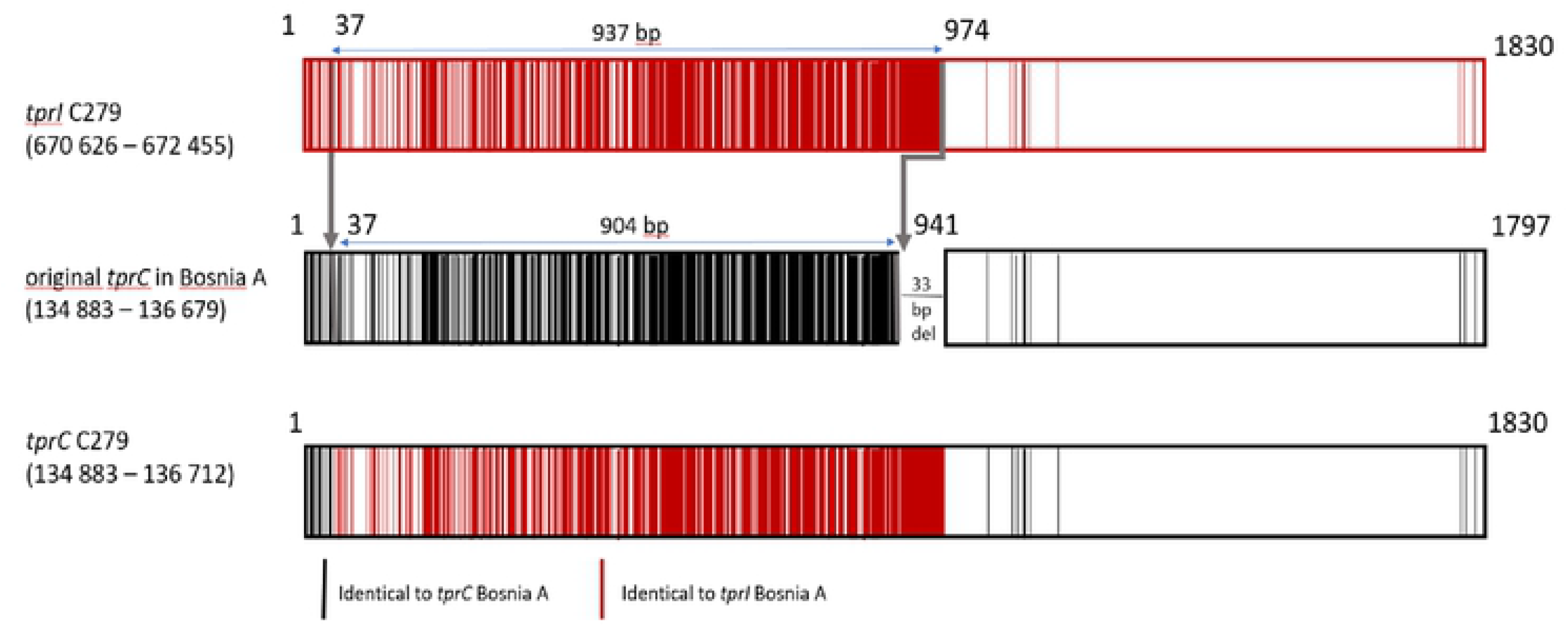
Overview of recombinant *tprC* in the C279 genome. Recombinant sites within the *tprC* gene and the resulting *tprCJ* chimera of C279. Black lines represent nucleotides identical to *tprC,* while red lines nucleotides from *tprl.* White color represents nucleotides shared between both *tprC* and *tprl* genes.

Between TP0127b and the pseudogene TP0128, the TEN genome C279 showed a 72 bp deletion followed by the duplication of a 53 bp-long sequence from TP0129. Similar to the *tprCI* chimera, the 53 bp-long sequence appears to be copied from TP0129 and inserted at the front of the TP0128 pseudogene. In other treponemal genomes, the region comprising TP0127-TP0129 was shown to be highly variable, including TPA Nichols, TEN Bosnia A, and TPeC strain Cuniculi A (Fig 3).

**Figure 3.**
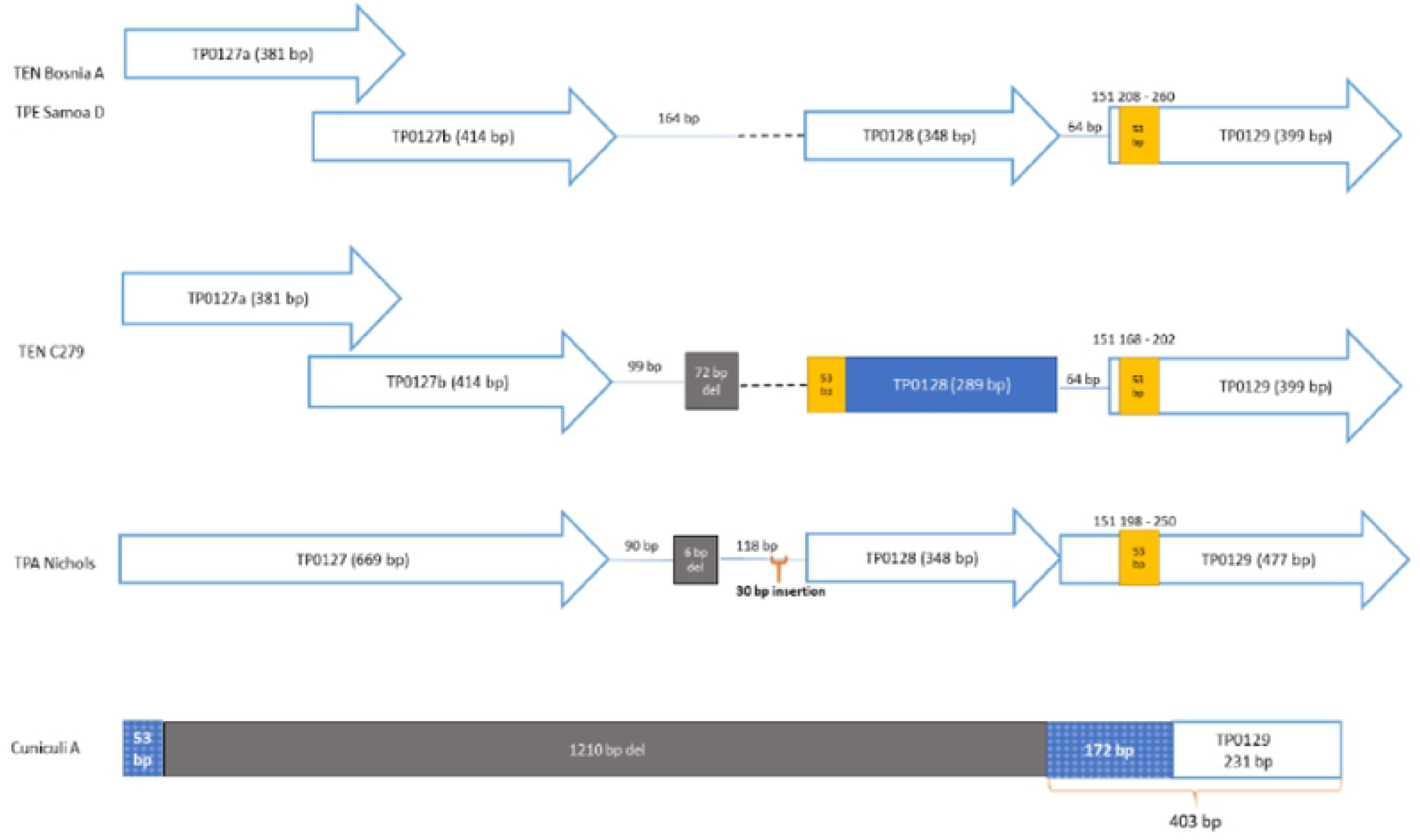
Comparison of region TP0127 TP0129 between TEN Bosnia A, TEN C279, TPA Nichols, TPE Samoa D, and TPeC Cuniculi A strain. TEN Bosnia A was used as the reference strain, and the TPE Samoa D strain was identical to TEN Bosnia A. The TEN genome C279 showed a 72 bp deletion and a 53 bp duplication compared to TP0129. TPA Nichols strain in IGR showed a 6 bp deletion and a 30 bp insertion compared to Bosnia A. Most of the sequence of Cuniculi A showed deletions, except for 225 bp, which was sequentially unrelated to TP0127-TP0128 in the other strains.

## Discussion

While bejel typically occurs in dry arid areas such as the Sahel and the Middle East – Cuba, with its hot, humid weather, is an atypical region for bejel to occur. In addition, Cuban bejel cases appear to be sexually transmitted, mostly among homosexual men, which is also atypical for bejel. While in several countries, including Canada and France, bejel cases were found to have been imported, no such indication exists for bejel in Cuba.

Whereas a previous paper on molecular typing of syphilis treponemes (TPA) revealed the presence of TEN, all of the nine independent isolates were found to be genetically indistinguishable, suggesting a clonal outbreak of imported cases of bejel [7,14]. A similar situation was recently published for Japanese bejel cases where it was found that all isolates were identical when two chromosomal loci were sequenced [6].

Since 2013, there have been numerous reports describing traces of genetic recombination events among pathogenic treponemes at more than a dozen genetic loci [32,33]. While there are examples of both intra- and inter-strain recombination events [15], in the genomes of the Cuban strains, we showed two new intra-strain recombination events. The intra-strain recombination event between *tprG* and *tprJ* has also been described in TPA, where a sequence from *tprJ* was copied to the *tprG* gene [15]. Here, we have described a new gene conversion event where the sequence from *tprI* was copied to the *tprC* locus. Previous studies also revealed changes at the TP0126–TP0127 loci in the TPA Nichols and TPE strains [34,35], showing that this region differs in gene prediction and region size. Moreover, the region comprising TP0129–TP0130 was previously described as one of the major donor sites for the variable regions of the *tprK* gene [36]. The emergence of the 53 bp-long sequences copied from the TP0129 by intra-strain gene conversion is therefore not surprising in this region.

The inter-strain recombination events in the TP0488 and TP0548 were described earlier [5]. From a previous study, we know that in most of the samples (C46, C75, C77, C131, C241, and C273), the TP0488 gene sequence was in the 11q/j isolate and therefore likely represented a recombinant allele [5]. Another similarity between C249 and isolate 11q/j was found in gene TP0865, which contains a 23 bp insertion in the same position as the 11q/j harbor 22 bp-long insertions. The size of insertions differs because of the different numbers of nucleotides in the homopolymeric tract. In both cases (i.e., in C249 and in 11q/j), the gene is considered to be non-functional, containing a frameshift mutation. In the C279 genome, the recombinant allele also seems to be present at the TP0548 locus, similar to Bosnia A and Iraq B versions but different from the TEN 11q/j strain [5].

Whole-genome sequencing revealed a surprising amount of genetic diversity within the Cuban isolates ranging between 0.2–10.3 differences per 100 kb, which corresponds to an estimated 22.8 to 117.4 nucleotide differences between individual strains on a genome-wide level. However, these changes are not visible in phylogenetic trees due to the differences in sequenced genome segments and omission of undetermined positions (N nucleotides) from the phylogenetic tree. This extent of genetic diversity is comparable to differences between TEN Bosnia and Iraq B with an average value of nucleotide diversity of 3.1, with 37 single-nucleotide differences, four indels, two differences in the number of tandem repetitions, and 18 differences in the length of homopolymeric regions were found in the Iraq B genome [12] compared to Bosnia A [37]. While TEN strain Iraq B was isolated in Iraq (the Middle East, southwestern Asia) in 1951, the strain Bosnia A was isolated in 1950 in Bosnia, southern Europe. A similar extent of genetic diversity was also found between TPE strains at different times and different geographical regions from which whole genome sequences are available (Samoa D, Gauthier, Kampung Dalan 363, Sei Geringging K403, CDC-1, CDC-2, CDC 2575, and Ghana-051) [35,38,39] ranging between 0.0–20.5 differences per 100 kb. This fact further supports the non-clonal character of Cuban TEN strains and is consistent with the long-term evolution of each of the Cuban TEN isolates. For comparison, TPA SS14-like strains obtained by direct sequencing [15] have nucleotide diversity in a range of 0.2–4.6.

Uncultivable pathogenic treponemes including TPA, TPE, and TEN are monomorphic bacteria [1,40], which have extremely high sequence similarity; therefore, it is likely that these related treponemes show similar mutation rates. While there are no studies on mutation rates in TEN, the mutation rate in TPE and TPA have been estimated as 1.21 × 10^−7^ and 0.82 × 10^−7^ per nucleotide site per year or lower, respectively [9,38]. Assuming that that mutation rate is similar in TEN as in TPA and TPE, the estimated 22.8 to 117.4 nucleotide differences between individual TEN strains detected in Cuba suggest at least several hundreds of years of separate evolution in the human population and, therefore, the long-term existence of the different TEN strains in the Cuban or related population.

Altogether, the findings presented in this study suggest that there are several different sequence types of TEN strains circulating in the population of Cuba. Evidence suggests that they are being transferred mainly through sexual transmission since all source patients were suspected of having syphilis, and most of them had ulcerations in the genital area. Unlike the established concept of treponemal subspecies and corresponding diseases, this study points to the fact that at least in the early stages of the disease, both bejel and syphilis treponemes produce symptoms that are indistinguishable. However, the role of the identified TEN genomes rearrangements in the development of similar symptoms remains unknown.

The authors recognize that the main limitation of the study was the low number of patients with high-quality samples suitable for sequencing; better samples would have allowed a more robust sequence analysis.

The non-clonal character of the Cuban TEN isolates suggests that Cuban TEN is a persistent infection within the human population rather than a single outbreak of a TEN isolate introduced from an area where it is typically endemic.

## Acknowledgments

We thank Thomas Secrest (Secrest Editing, Ltd.) for the English editing of the manuscript.

## Author Contributions

Data curation: Eliška Vrbová

Formal analysis: Eliška Vrbová, Angel A. Noda, Linda Grillová, Allyn Forsyth, Jan Oppelt, David Šmajs

Funding acquisition: David Šmajs

Investigation: Eliška Vrbová, Angel A. Noda, Linda Grillová, Islay Rodríguez, Allyn Forsyth, Jan Oppelt, David Šmajs

Methodology: Eliška Vrbová, Angel A. Noda, Linda Grillová, , Allyn Forsyth, Jan Oppelt

Supervision: David Šmajs

Validation: Linda Grillová, Allyn Forsyth, Jan Oppelt, David Šmajs

Writing – original draft: Eliška Vrbová, Angel A. Noda, David Šmajs

Writing – review & editing: Eliška Vrbová, Angel A. Noda, Linda Grillová, Islay Rodríguez, Allyn Forsyth, Jan Oppelt, David Šma

## Supplementary data

Supplementary data are available.

